# Overinflation and overconcentration: why Cauchy perturbation kernels are the right choice for ABC-SMC

**DOI:** 10.64898/2026.06.24.734205

**Authors:** Marc Sturrock, Vahid Shahrezaei

## Abstract

Approximate Bayesian computation sequential Monte Carlo (ABC-SMC) propagates its particles with a perturbation kernel, and with the standard Normal kernel it degrades sharply as the parameter dimension grows, a failure usually attributed to dimension itself. We show instead that it is governed by the quality of the summary statistics, with dimension entering only through a separate and milder mechanism, and that the two must act together for the Normal kernel to break. The first ingredient is covariance overinflation: the kernel covariance, estimated from the particle cloud, overshoots the true posterior covariance by a factor set by information loss in the summary statistics. We derive this overscaling factor in closed form for a Gaussian model with sufficient statistics and show that it stays modest at any dimension, shrinking toward its baseline value as the tolerance tightens; the extreme values seen in practice (of order 10^3^) are a signature of insufficient summaries, not of dimension. The second ingredient is perturbation overconcentration: the normalised Normal step size concentrates around one as the dimension grows, so every proposal overshoots by the same factor. Either ingredient alone is harmless; only their combination breaks the Normal kernel. A Cauchy kernel (multivariate *t* with one degree of freedom) removes the concentration, keeping a positive acceptance rate under arbitrary overscaling at a bounded worst-case cost of 1.87× in expected squared jump distance. In a Metropolis–Hastings framework we derive closed-form acceptance rates for both kernels that illustrate the advantage of the Cauchy kernel in this limit. A series of full ABC-SMC computational experiments on five problems at *d* = 12, including a hierarchical gene-expression model, show the Cauchy reducing the sliced Wasserstein distance to the reference posterior by factors of up to 50 with the same simulation budget. Since the summary statistics are commonly insufficient for the models that require ABC, overinflation is structural and the Cauchy perturbation kernel is the right default for problems in higher dimensions.

## 1 Introduction

Approximate Bayesian computation (ABC) enables posterior inference for models with intractable likelihoods by replacing likelihood evaluation with forward simulation and comparison of summary statistics (Sisson et al., 2007; Toni et al., 2009; Beaumont et al., 2009). Sequential Monte Carlo (SMC) variants of ABC propagate a population of weighted particles through a sequence of decreasing tolerance thresholds, using a perturbation kernel to propose new parameter values at each iteration. Throughout this paper we use “Normal” when referring to the perturbation kernel and “Gaussian” when referring to the target model, to avoid ambiguity when both appear together. When the dimension of the parameter space (*d*) exceeds about five, Normal perturbation kernels degrade rapidly: acceptance rates collapse, the algorithm stalls, and the ABC posterior fails to concentrate on the true posterior. Replacing the Normal kernel with a Cauchy kernel (multivariate *t* with one degree of freedom) recovers much of the lost performance. This Cauchy advantage has been attributed to “high dimension,” but that attribution conflates two distinct mechanisms that happen to worsen together as dimension grows.

Research on kernel choice in ABC-SMC has focused primarily on adaptation, learning the scale and orientation of the kernel from the particle population, rather than on the shape of the kernel, i.e. its tail behaviour. Beaumont et al. (2009) introduced the now-standard practice of setting the kernel covariance to twice the weighted empirical covariance. Filippi et al. (2013) derived optimality criteria based on minimising the Kullback–Leibler divergence between successive ABC target distributions, but restricted attention to Gaussian kernel families and locally adapted variants. Prangle et al. (2025) recently conducted an extensive empirical comparison of ABC-SMC kernels and recommend a one-hit kernel with a Gaussian mixture proposal. Every kernel they consider is nonetheless Gaussian-based, their benchmarks remain low-dimensional (their largest example has *d* = 5), and they provide no mechanism for the performance differences they observe. Heavy-tailed perturbations are left unexamined, and a Gaussian mixture, however flexible, retains Gaussian tails, so each of its components still concentrates onto a thin shell as *d* grows. The question of whether the Gaussian shape is appropriate when the kernel covariance is structurally misestimated has not been addressed.

The MCMC literature provides theoretical grounding for the fragility of Gaussian proposals. Roberts et al. (1997) derived the celebrated 0.234 optimal acceptance rate for Gaussian random-walk Metropolis and showed that the proposal variance must scale as *O*(*d*^−1^), making performance exquisitely sensitive to the scale parameter. Roberts and Rosenthal (2001) formalised optimal scaling across Metropolis–Hastings variants, and Sherlock and Roberts (2009) extended the analysis to elliptically symmetric targets, showing that the probability mass mass of a Gaussian perturbation concentrates in a thin hyperspherical shell whose radius grows as 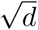 . Sherlock et al. (2010) synthesised these results, highlighting the practical consequence: when the proposal scale is even modestly wrong, the entire shell of proposed moves misses the target, and the chain stalls. In ABC-SMC the proposal scale is never correctly specified; it is estimated from a noisy, weighted particle cloud, so this fragility is not a tuning failure but a structural one.

Independently, the global optimisation community discovered the same heavy-tailed advantage from a different direction. Yao et al. (1999) showed that replacing Gaussian mutation with Cauchy mutation in evolutionary programming dramatically improves performance on multi-modal benchmark functions, because the Cauchy’s long tails permit occasional large jumps that escape local optima. Earlier, Szu and Hartley (1987) proved that Cauchy-distributed state perturbations in simulated annealing permit an inverse-linear cooling schedule *T* (*k*) ∝ 1*/*(1 + *k*), exponentially faster than the inverse-logarithmic schedule required by Gaussian perturbations. This result has a direct analogue in ABC-SMC: the Cauchy kernel permits more aggressive tolerance reduction without freezing the particle population, because its variable step sizes ensure that some fraction of proposals remain close to the current position even when the kernel covariance is overscaled. The Lévy flight literature formalises the underlying principle: in sparse search environments, heavy-tailed step-length distributions with power-law exponent *α* = 1 (the Cauchy case) optimise the rate of discovering new targets (Viswanathan et al., 1999). High-dimensional ABC, where the acceptance region occupies a vanishing fraction of the prior volume, is precisely such a sparse environment.

The weight degeneracy that accompanies high-dimensional SMC is well understood in the particle filtering literature; we examine it directly for both perturbation kernels in Section 3.5. Snyder et al. (2008) showed that the ensemble size required to avoid filter collapse grows exponentially with the variance of the log-likelihood increments, and Bengtsson et al. (2008) proved a sharp phase transition: below a critical particle count the filter collapses entirely. Del Moral et al. (2006) analysed weight variance in general SMC samplers and established conditions under which adaptive proposals can control degeneracy. In ABC-SMC, the kernel covariance estimated from a degenerating particle cloud feeds directly back into the proposal, creating a vicious cycle: overinflated covariance → concentrated Gaussian shell misses target → weight collapse → worse covariance estimate. This paper identifies covariance overinflation and step-size concentration as the two components of this cycle and shows that the Cauchy kernel breaks it by eliminating the concentration step.

Specifically, we disentangle two independent failure modes. The first is covariance over-inflation: the kernel covariance overshoots the true posterior covariance by a large factor 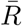 because the summary statistics lose information. In a Gaussian model with sufficient statistics, 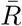 is modest at any dimension; the extreme values observed in practice 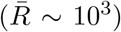 are due to insufficient summary statistics. The second is perturbation overconcentration: at high *d*, the normalised Normal step size *χ*^2^(*d*)*/d* concentrates around 1 with standard deviation 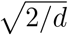, so every proposal overshoots by exactly the same factor. Overinflation without concentration is harmless (at *d* = 3 with 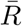 = 1,600 both kernels perform identically), and concentration without overinflation is harmless (with sufficient statistics 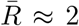 and the Normal kernel works well). Both are necessary for the Normal kernel to fail; removing either one saves it. Unlike prior work on kernel optimality, which assumes the covariance is well estimated, we show that the interaction of structural covariance overinflation with Gaussian step-size concentration is the root cause of ABC-SMC failure in moderate dimensions.

Section 2 describes the test problems and evaluation metrics. Section 3 presents the results: documenting both failure modes empirically, deriving the overinflation ratio analytically, proving that the Cauchy acceptance rate stays positive under arbitrary overscaling, verifying the predictions, and comparing the two kernels in full ABC-SMC. Section 4 discusses the implications and makes the case for the Cauchy as the default perturbation kernel in ABC-SMC.

## 2 Methods

We use five test problems that span a range of posterior geometries and summary statistic quality, from exact sufficiency to severe information loss. All are parameterised by dimension *d* and tested at *d* = 3 and *d* = 12.

### 2.1 Gaussian

The simplest problem: *X*_*i*_ ∼ *N* (*θ, I*_*d*_) for *i* = 1, …, *n* with *n* = 50 and a flat prior on [−10, 10]^*d*^. The true posterior is 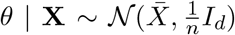 . The summary statistic 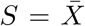 is sufficient, so this problem serves as a baseline where overinflation should be minimal. Reference posterior samples are drawn analytically.

### 2.2 Neal funnel

This problem is an ABC variant of the funnel construction of Neal (2003). The likelihood couples a scalar parameter *θ*_1_ to a (*d* − 1)-dimensional location *θ*_2:*d*_ through a heteroscedastic noise model:

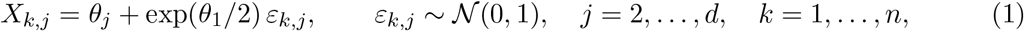

with *n* = 100. Priors are uniform: *θ*_1_ ∼ *U* (−9, 9) and *θ*_*j*_ ∼ *U* (−15, 15) for *j* = 2, …, *d*, matching ±3*σ* of the Normal priors typically used for this target. Although the priors are flat, the posterior inherits the classical funnel geometry from the likelihood: the conditional spread of *θ*_2:*d*_ varies by orders of magnitude with *θ*_1_ through the exp(*θ*_1_*/*2) noise scale. The summary statistics are the component wise sample means and variances, and the ABC distance is 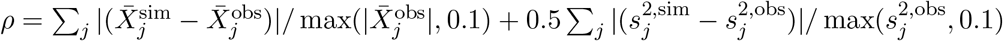 .

The posterior is analytically tractable. Conditional on *θ*_1_, the remaining parameters are independent Gaussians with known mean and variance (depending on *θ*_1_ and the data), so we can compute the marginal posterior of *θ*_1_ on a grid and sample the conditionals exactly. Reference samples and the true posterior covariance are computed from 10^5^ analytical samples.

### 2.3 Banana

The Banana problem has *d* + 1 parameters *θ* = (*θ*_1_, …, *θ*_*d*+1_): the first *d* are a location *x* = (*θ*_1_, …, *θ*_*d*_) on the curved ridge of the Rosenbrock function

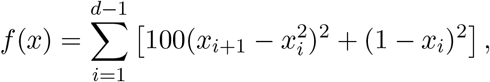

and the last, *θ*_*d*+1_, is a log-noise level. A dataset consists of *n* = 50 scalar measurements of the Rosenbrock surface, each corrupted independently by heteroscedastic Gaussian noise:

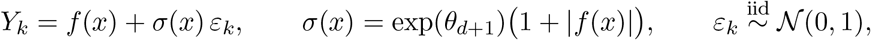

for *k* = 1, …, *n*; equivalently *Y*_*k*_ ∼ *N* (*f* (*x*), *σ*(*x*)^2^) . The location *x* thus enters both through the signal *f* (*x*) and through the noise scale, whereas *θ*_*d*+1_ controls the overall noise level only.

The summary statistics are quantiles of the simulated outputs at 19 equally spaced levels from the 5th to the 95th percentile. The posterior concentrates on a thin curved ridge whose geometry is not well captured by a Gaussian covariance, making this a worst case for covariance-based methods. Reference posterior samples were obtained from the No-U-Turn Sampler (NUTS; Hoffman and Gelman, 2014) with 10^5^ post-warmup draws.

### 2.4 Shell

The Shell problem has *d* parameters constrained to lie near a hypersphere of radius *R* = 3 in ℝ^*d*^. Observations are noisy projected radii: a dataset consists of *n* = 100 values *Y*_*k*_ = ∥*θ* + *σ*_*w*_ *ε*_*k*_∥, where 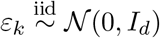 is a standard Gaussian perturbation of the parameter vector *θ* and *σ*_*w*_ = 0.3 sets its scale.

Summary statistics are the mean and standard deviation of the projected radii. The posterior is annular, with no density at the origin (the mode of the prior), making it a test of non-convex posterior geometry. Reference posterior samples were obtained from NUTS with 10^5^ post-warmup draws.

### 2.5 Gene expression

The preceding test problems are synthetic. To demonstrate the Cauchy kernel advantage on a problem grounded in biology, we include a hierarchical gene expression model motivated by single-cell RNA sequencing (scRNA-seq).

The telegraph model of gene expression describes a gene that switches between inactive and active states, with mRNA produced only in the active state (Peccoud and Ycart, 1995; Shahrezaei and Swain, 2008):

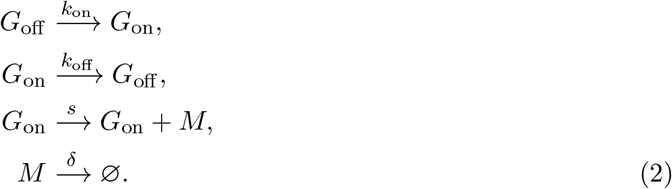

In the bursty limit (*k*_off_ ≫ *k*_on_, *s* ≫ *k*_off_), transcription occurs in short bursts of geometrically distributed size and the steady-state mRNA count follows a Negative Binomial distribution with mean *μ* = *k*_on_*s/*(*k*_off_ *δ*) and dispersion *ϕ* = *k*_on_*/δ*. We adopt a hierarchical Gamma-Poisson formulation in which *d* genes share a common dispersion parameter *ϕ* but have gene-specific mean expression levels drawn from a log-normal population distribution:

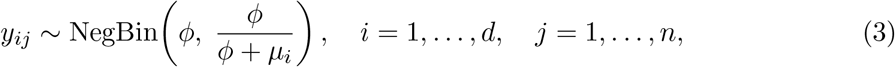

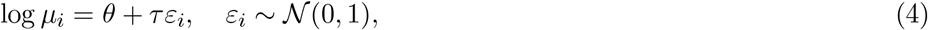

where *θ* is the global log-mean expression, *τ* controls the spread of gene-specific means, and *ϕ* is the shared overdispersion. This structure is standard in scRNA-seq analysis: tools such as BASiCS (Vallejos et al., 2015), scran (Lun et al., 2016), and DESeq2 (Love et al., 2014) all employ hierarchical shrinkage of gene-specific parameters toward a shared population distribution. The data consist of a single snapshot of *n* = 100 cells across *d* genes. The negative-binomial count model is empirically validated as the scRNA-seq noise model by Grün et al. (2014), and the log-normal distribution of gene-specific expression levels was demonstrated empirically for single-cell mRNA by Bengtsson et al. (2005).

The parameter vector is (*θ, τ, ϕ, ε*_1_, …, *ε*_*d*_), giving *d* + 3 parameters. The summary statistics are the analytical per-gene mean and variance (2*d* statistics):

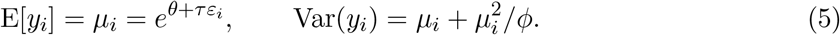

The ABC distance compares these analytical moments with observed sample moments: 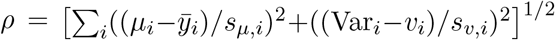, where *s*_*μ,i*_ and *s*_*v,i*_ are componentwise scales. Because the moments are deterministic functions of the parameters, each ABC distance evaluation is instantaneous (no forward simulation required).

Priors are uniform, as is standard in ABC applications: *θ* ∼ *U* (−4, 8), *τ* ∼ *U* (0.001, 10), *ϕ* ∼ *U* (0.1, 50), and *ε*_*i*_ ∼ *U* (−3, 3). Reference posterior samples were obtained from NUTS with 2.4 × 10^4^ post-warmup draws using the full Negative Binomial likelihood on all *d* × *n* counts (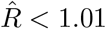 for all parameters).

### 2.6 Evaluation metrics

For each problem, the true posterior covariance Σ was estimated from reference samples (analytical or NUTS).

Posterior quality is assessed by three variants of the classifier two-sample test (C2ST; Lopez-Paz and Oquab, 2017), each comparing resampled ABC posterior samples against reference samples using a neural network classifier with 5-fold cross-validation. The joint C2ST trains the classifier on all *d* dimensions simultaneously and is the most powerful test. The marginal C2ST (mC2ST) averages *d* separate one-dimensional classifiers, detecting per-margin differences but missing correlation structure. The pairwise C2ST (pC2ST) averages 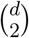 classifiers on all pairs of dimensions, capturing pairwise correlations without the curse of dimensionality of the full joint test. For all three, 0.5 means the ABC and reference samples are indistinguishable and 1.0 means they are perfectly separable. However, the joint C2ST saturates at 1.0 for most *d* = 12 cases, limiting its ability to discriminate between methods in higher dimensions.

To complement the C2ST variants we report the sliced Wasserstein distance (SWD; Bonneel et al., 2015), a proper metric between distributions that does not saturate. The SWD projects both sample sets onto *L* = 1,000 random unit vectors ***θ*** ∈ *S*^*d*−1^, computes the one-dimensional Wasserstein-1 distance (via sorted samples) along each projection, and averages:

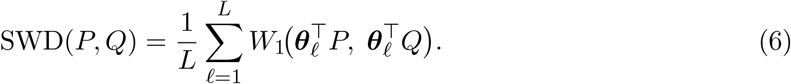

Lower values indicate closer agreement with the reference posterior; SWD = 0 if and only if the distributions are identical. Unlike C2ST, SWD provides a continuous distance that scales well to moderate dimensions and retains dynamic range where the classifier-based tests saturate.

We also report the effective sample size (ESS) of the importance weights, a standard diagnostic for weight degeneracy. For weights *w*_1_, …, *w*_*N*_ at the final ABC-SMC iteration,

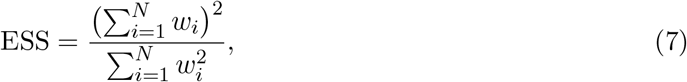

which equals *N* when the weights are uniform and falls toward 1 as a few particles dominate. We report it as the fraction ESS*/N* ∈ (0, 1].

Unless otherwise stated, all ABC-SMC runs use *N* = 4,000 particles with the standard Toni et al. (2009) prescription: the kernel covariance is set to twice the weighted empirical covariance at each iteration, and the tolerance is reduced to the median distance at each step.

## 3 Results

### 3.1 ABC-SMC produces overinflated kernel covariances

To quantify the mismatch between the scale matrix 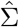 used by the perturbation kernel (twice the weighted empirical covariance of the current particle cloud) and the true posterior covariance Σ, we define the mean overscaling ratio

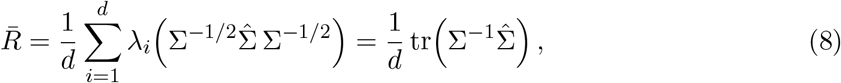

where *λ*_*i*_(·) denotes the *i*-th eigenvalue, so 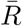 is the mean of the eigenvalues of the kernel scale matrix taken relative to the posterior. The Normal and Cauchy kernels are both parameterized by this same 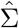 (for the Normal kernel it is the covariance; for the heavy-tailed Cauchy, which has no covariance, it is a scale matrix), so 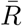 depends only on the two matrices and is computed identically for either kernel, independently of the kernel’s tail shape. The functional 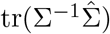, a scale-invariant comparison of two positive-definite matrices, is the trace term of Stein’s loss for covariance estimation (James and Stein, 1961; Dey and Srinivasan, 1985); 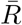 is its normalisation by the dimension *d*.

At 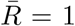 the kernel matches the posterior; at 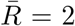 it is twice as wide in every direction (the minimum achievable under the standard 2× covariance prescription). The true posterior covariance Σ is estimated from reference samples: either analytical draws (Gaussian, Funnel) or NUTS output (Banana, Shell).

We ran vanilla ABC-SMC on the Gaussian model (sufficient statistics) and the Funnel (insufficient summary statistics) at *d* ∈{3, 7, 12} with *N* = 4,000 particles, a simulation budget of 10^8^, and *p*_acc,min_ = 10^−4^, averaging 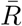 trajectories over three independent replicates.

Figure 1 contrasts the two problems. On the Gaussian (left panel), all three trajectories decline toward 2. At *d* = 3 and *d* = 7, 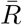 converges to the prescription factor within 26 and 40 iterations respectively, confirming that overinflation can be eliminated when summaries retain full information. At *d* = 12 the algorithm reaches 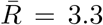 after 53 iterations and is still declining. On the Funnel (right panel), all trajectories trend toward 2, but convergence is dramatically slower. At *d* = 3, 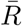 reaches 2 after 29 iterations; at *d* = 7 it is still at 5 after 43 iterations; and at *d* = 12 it remains above 150 after 56 iterations despite consuming the full 10^8^ simulation budget. The overinflation is not permanent, but insufficient summary statistics make convergence intractably slow, particularly in higher dimensions.

**Figure 1:**
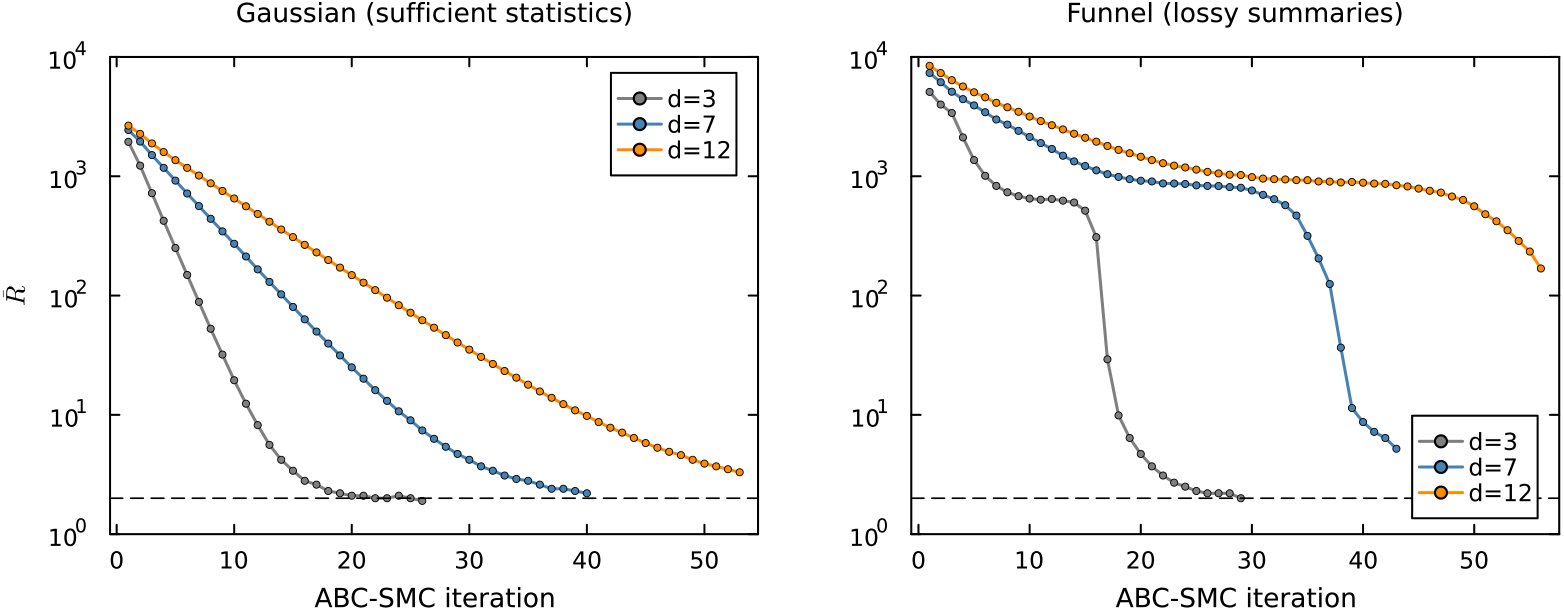
Mean overscaling ratio 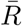 across ABC-SMC iterations using a Normal perturbation kernel (*N* = 4,000, 10^8^ simulations, *p*_acc,min_ = 10^−4^, averaged over 3 replicates). Left: Gaussian model with sufficient statistics; all dimensions converge toward 2 (dashed line). Right: Funnel with insufficient summary statistics; all trajectories trend toward 2 but convergence becomes intractably slow with increasing dimension.

This overscaling is structural: it is a property of the ABC-SMC construction itself, not a tuning mistake or a finite-sample artefact. At each iteration the kernel covariance is set to twice the weighted empirical covariance of the particle population, so the standard prescription already places a baseline of 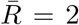, the value reached if the population perfectly matched the target. The population, however, approximates the ABC posterior *π*_*ϵ*_(*θ*) ∝ *π*(*θ*) Pr(*ρ < ϵ*_*t*_ | *θ*), which is wider than the true posterior whenever *ϵ*_*t*_ *>* 0. The kernel covariance inherits this excess width, so in practice 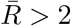, with the gap above 2 governed by how far the current tolerance *ϵ*_*t*_ is from zero. As *ϵ*_*t*_ → 0 the gap closes and 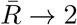 from above; how quickly depends on the problem: fast when the summaries are informative and *d* is small (Figure 1, left), intractably slow otherwise (right panel). Section 3.3 analyses what sets the size of this gap; here we establish only that it does not shrink with the particle count.

One might hope that increasing the particle count would reduce overscaling. Table 1 shows it does not. We ran the Funnel at *d* = 12 with *N* ranging from 500 to 8,000 using a Normal kernel. The final 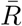 is flat at roughly 870 across all particle counts, varying by less than 4%. The overscaling is not a sample-size artefact; it is a convergence property of the algorithm at this dimension.

**Table 1:** Final overscaling ratio 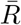 across particle counts (Funnel, *d* = 12, Normal kernel, *p*_acc,min_ = 0.01, 10^8^ sim budget, 3 replicates per row; the reported values are 3-rep means). 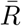 does not decrease as *N* grows from 500 to 8,000, indicating that overscaling is structural rather than a sample-size limitation. ESS/*N* : effective sample size as a fraction of the particle count at the final iteration.

| $N$ | Iterations | $\bar{R}$ | Total sims | ESS/$N$ |
| --- | --- | --- | --- | --- |
| 500 | 40 | 852 | 1.8M | 81% |
| 1,000 | 40 | 863 | 3.6M | 78% |
| 2,000 | 39 | 872 | 7.0M | 81% |
| 4,000 | 40 | 878 | 14.1M | 82% |
| 8,000 | 40 | 880 | 28.5M | 81% |

### 3.2 Normal perturbation kernels overconcentrate at high dimensions

The second ingredient is perturbation overconcentration: at high *d*, every Normal proposal lands at the same Mahalanobis distance from the current point.

A Normal perturbation with covariance *k*Σ has squared Mahalanobis step size *k* · *χ*^2^(*d*). The normalised step *χ*^2^(*d*)*/d* has mean 1 and standard deviation 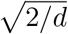 :

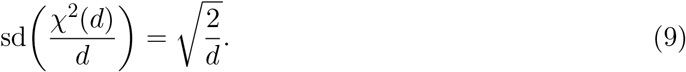

At *d* = 3 this is 0.82; at *d* = 12 it is 0.41; at *d* = 100 it is 0.14. The step size concentrates onto a thin shell of squared radius *kd*, and the shell becomes thinner with increasing *d*.

A multivariate Cauchy (*t*_1_) perturbation is a Normal perturbation scaled by a random factor 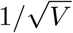, where *V* ∼ *χ*^2^(1) is drawn independently for each proposal. If *V* were constant, this would simply rescale the Normal and the concentration problem would remain. But *V* is random with coefficient of variation 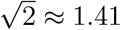 regardless of *d*: when *V* is large, the step is much shorter than the Normal and can land inside the acceptance region; when *V* is near zero, the step overshoots by orders of magnitude.

Figure 2 shows the physical perturbation distance ∥*ε*∥ for both kernels across three dimensions (*d* = 3, 12, 20) and three levels of kernel covariance overinflation 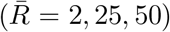. Reading down the rows, increasing dimension shifts both distributions rightward and narrows the Normal into a spike: at high *d* every Normal perturbation lands at nearly the same distance, with none short enough to be useful. The Cauchy is also affected by dimension (it too loses mass near zero as *d* grows) but retains a wider spread than the Normal at every *d*. Reading across the columns, increasing overinflation stretches both distributions further from the origin, and the Cauchy stretches asymmetrically, holding a longer left tail than the Normal. That asymmetry is the source of its advantage: some Cauchy proposals remain short enough to land in the acceptance region even when the kernel covariance overshoots the posterior.

**Figure 2:**
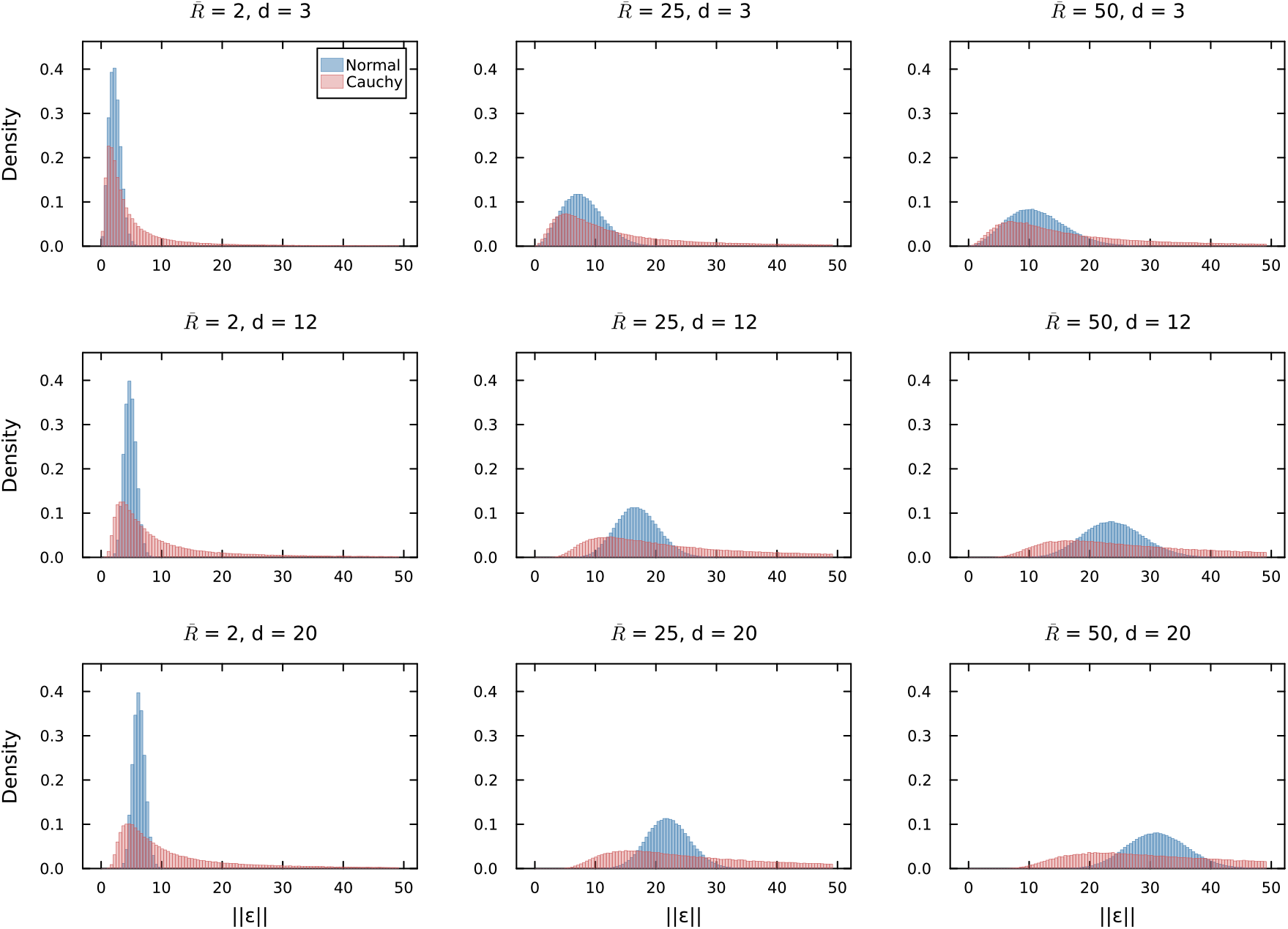
Physical perturbation distance ∥*ε*∥ from 200,000 random draws for Normal (blue) and Cauchy (red) kernels. Rows: *d* = 3 (top), *d* = 12 (middle), *d* = 20 (bottom). Columns: kernel covariance overinflation 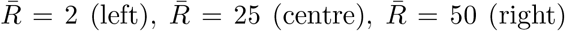 . Increasing dimension shifts both distributions rightward and narrows the Normal into a spike. Increasing overinflation stretches both distributions, but the Cauchy retains a longer left tail. All panels share the same axis scales.

These effects are properties of the proposal distribution alone; they do not depend on the target, the summary statistics, or the ABC tolerance. Dimension removes the possibility of short perturbations for both kernels; overinflation determines how short a perturbation needs to be for acceptance. The Cauchy advantage arises when both are present: it cannot eliminate the dimensional floor on step size, but it can produce perturbations that are shorter than the Normal’s by a factor governed by 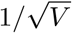 .

### 3.3 Overinflation is driven by summary statistic quality, not necessarily dimension

Consider a *d*-dimensional Gaussian model: *X*_*i*_ ∼ *N* (*θ, σ*^2^*I*_*d*_) for *i* = 1, …, *n*, with *σ*^2^ known and a flat prior on *θ*. The sufficient statistic is 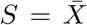, and the true posterior is *θ* |*S*_obs_ ∼ *N* (*S*_obs_, (*σ*^2^*/n*) *I*_*d*_).

ABC-SMC uses the distance *ρ* = ∥*S*_sim_ − *S*_obs_∥ and accepts when *ρ < ϵ*. Since *S*_sim_ | *θ* ∼ *N* (*θ*, (*σ*^2^*/n*) *I*_*d*_), the acceptance probability at parameter value *θ* is

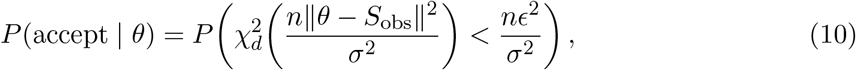

where 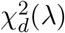 denotes a non-central chi-squared random variable with *d* degrees of freedom and non-centrality *λ*.

Define the normalised tolerance *η* = *ϵ/σ*_post_ where 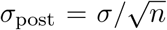 is the posterior standard deviation per direction. In normalised coordinates *u* = (*θ* − *S*_obs_)*/σ*_post_, the true posterior has unit variance per direction and the ABC posterior has radial profile

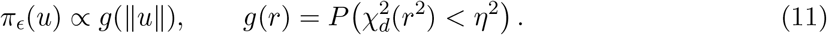

The mean overscaling ratio is

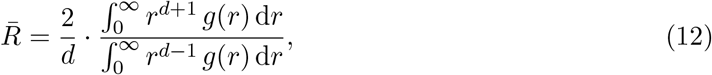

where the factor of 2 is the standard kernel covariance prescription 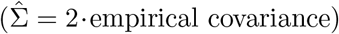 .

Equation (12) depends only on *η* and *d*. For tolerances much larger than the posterior scale, *g*(*r*) ≈ 1 throughout *r < η* and the integrals reduce to moments of a uniform ball, yielding

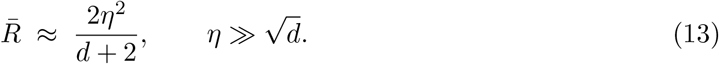

Figure 3 shows the acceptance rate at *θ* = *θ*_true_ as a function of normalised tolerance 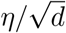, computed by numerical integration of (12) for *d* ∈ {3, 7, 12}. The dashed line marks 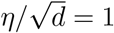 . At this point, the acceptance rate is 55–61% and 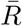 ranges from 3.2 to 3.7 (Table 2). Even at 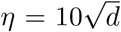, 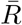 is only 122–173. To reach the empirical values of 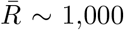 observed on the Funnel, the normalised tolerance would need to be 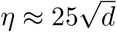, far above 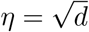 .

**Table 2:** Overinflation 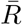 and acceptance rate (in parentheses) in the Gaussian model at selected normalised tolerances.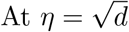, 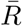 is between 3 and 4 regardless of dimension.

| $\eta/\sqrt{d}$ | $d = 3$ | $d = 7$ | $d = 12$ |
| --- | --- | --- | --- |
| 1 | 3.2 (61%) | 3.6 (57%) | 3.7 (55%) |
| 2 | 6.8 (98%) | 8.2 (98%) | 8.9 (99%) |
| 10 | 122 (100%) | 158 (100%) | 173 (100%) |

**Figure 3:**
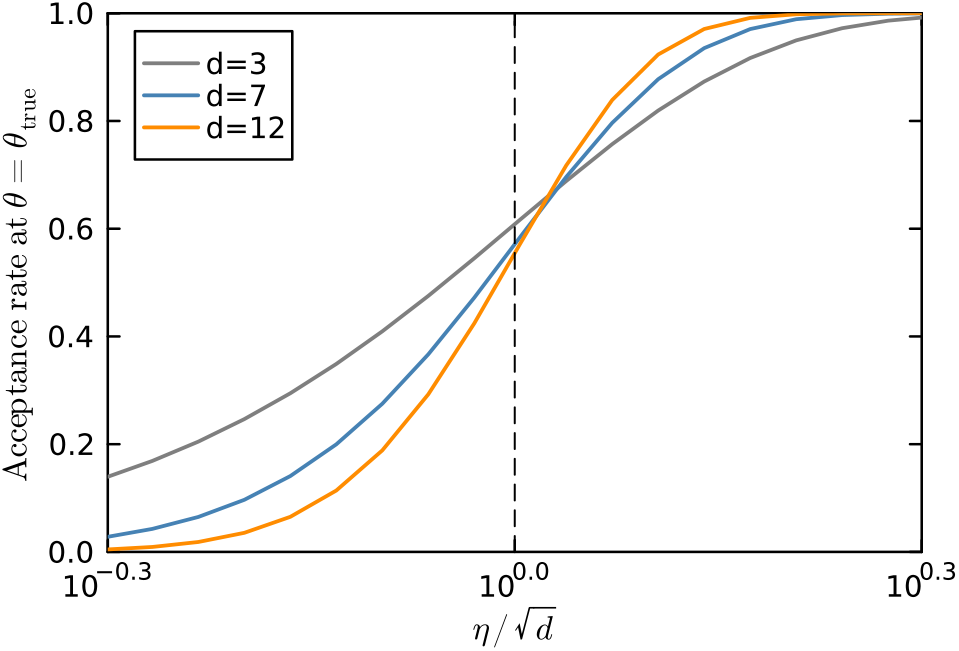
ABC acceptance probability at *θ* = *θ*_true_ in the Gaussian model with sufficient statistics, plotted against the tolerance *η* = *ϵ/σ*_post_ normalised by 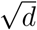 . The dashed line marks 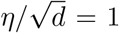, where 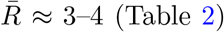 (Table 2). Normalising by 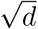 approximately collapses the curves, showing that the acceptance rate depends primarily on tolerance relative to the posterior scale, with only a modest residual effect of dimension.

Table 2 gives representative values.

With sufficient statistics, the ABC posterior converges to the true posterior as *ϵ* → 0, and 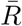 converges to 2 (the prescription factor) at any dimension. The extreme overinflation observed in practice requires insufficient summary statistics: a gap between the ABC posterior and the true posterior that does not close as the tolerance decreases.

To confirm that summary statistic quality, not dimension, drives overinflation, we ran ABC-SMC on the Gaussian model at *d* = 3 with deliberately degraded summaries. The model is *X*_*i*_ ∼ *N* (*θ, I*_3_) with *n* = 50 observations, so the true posterior has per-direction variance 1*/*50 = 0.02. We tested five summary statistic configurations, from sufficient to severely insufficient, each with *N* = 4,000 particles and 5 replicates.

The results in Table 3 confirm the analytical prediction. With sufficient statistics, 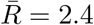, close to the analytical value of 3.2 at 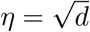 (the small gap arises because the algorithm pushes *η* slightly below 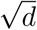). Once Gaussian noise with standard deviation 5 is added to that sufficient statistic, 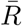 climbs above 1,600 at *d* = 3, putting the controlled-degradation experiment in the same regime as the Funnel at *d* = 12 ( 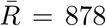, Table 1). Collapsing the three-dimensional summary to a scalar norm gives 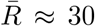, a fifteenfold increase. By holding dimension fixed and varying only summary quality, the experiment isolates one driver of overinflation. Cross-problem comparisons of 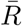 (Table 7) conflate that driver with model structure, since each problem necessarily uses different summary statistics and “equivalent” summaries are not well defined across models.

**Table 3:**
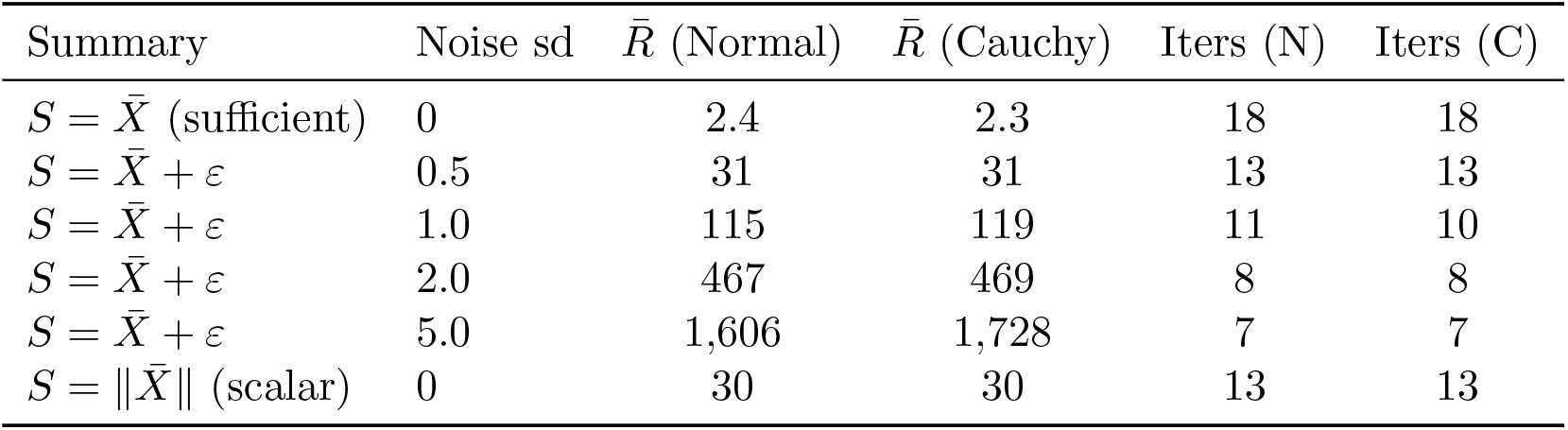
Overinflation at *d* = 3 under degraded summary statistics (Gaussian model, *N* = 4,000). Adding noise to the sufficient statistic 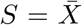 drives 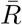 from 2.4 to over 1,600 without changing the dimension. Both kernels see the same 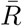 and complete the same number of iterations.

This separation matters because overinflation and overconcentration are independent quantities: the former is set by summary quality, the latter by dimension, and neither in isolation breaks the Normal kernel. Three regimes make the point.

When overinflation is severe but the dimension is low, the Normal kernel still works. At *d* = 3 with 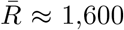 (Table 3), both kernels complete the same number of iterations (seven) and reach the same final 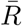 . Since sd(*χ*^2^(3)*/*3) = 0.82, the Normal step retains enough variability for some proposals to land close enough to be accepted, even when the typical step overshoots by a factor of 1,600.

When the dimension is high but overinflation is mild, the Normal kernel also works. The Gaussian model with sufficient statistics at *d* = 12 has 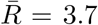 at 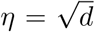 (Table 2). This is close enough to optimal scaling that the thin-shell effect is harmless: the shell at squared radius ≈ 3.7*d* sits within a small multiple of the acceptance region at squared radius ≈ *d*, and the Normal is 1.87× faster than the Cauchy in expected squared jump distance.

Only when both factors bite at once does the Normal collapse. On the Funnel at *d* = 12 with insufficient summary statistics, 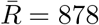 (Table 1) and sd(*χ*^2^(12)*/*12) = 0.41, so the Normal step is locked onto a shell at squared radius ≈ 878 · 12 = 10,536 while the acceptance region has squared radius ≈ 12. No Normal proposal can fluctuate three orders of magnitude inwards; the Cauchy’s 1*/V* factor supplies precisely that fluctuation.

In practice the two drivers tend to worsen together with dimension, which is why the Cauchy advantage can look dimension-driven from a distance. Summary statistics typically lose more information as *d* grows, because the number of moment-based statistics scales linearly with *d* but the complexity of the posterior need not. Problems like the Funnel possess conditional independence structure that marginal moments cannot capture, and the mismatch widens at higher *d*. Overconcentration, by contrast, is a monotone function of *d* through equation (9); no analogous monotone link runs from *d* to overinflation, since the Gaussian model with sufficient statistics shows 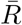 nearly constant across *d*.

Separating the two effects clarifies when a heavy-tailed kernel is actually needed. It is not high dimension per se that demands the Cauchy, but the combination of insufficient summary statistics and enough dimensions for concentration to bite. A three-dimensional problem with poor summaries will have a high 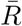 but gain nothing from the Cauchy; a twenty-dimensional problem with near-sufficient summaries will have a low 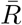 and gain nothing either. The Cauchy is the right default precisely when both conditions hold.

### 3.4 Cauchy perturbation kernel acceptance rates remain above zero under arbitrary overscaling

Switching from a Normal to a Cauchy kernel in ABC-SMC changes two things at once: the proposal distribution and the importance weights (since the kernel density appears in the weight denominator). To isolate the proposal effect, we work in the Metropolis–Hastings (MH) framework, where the kernel is used only to propose and there are no importance weights. If the Cauchy advantage survives in this cleaner setting, it must be a property of the proposal mechanism itself.

Let Σ be the true target covariance and suppose the proposal covariance is 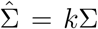 with *k >* 1. A Normal perturbation is Δ_N_ ∼ *N* (0, *k*Σ). A Cauchy perturbation (multivariate *t* with *v* = 1) can be written as

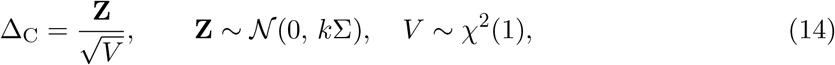

with **Z** and *V* independent. A Cauchy proposal is a Normal proposal scaled by a single random number 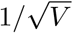, drawn once per proposal. The squared Mahalanobis step sizes are

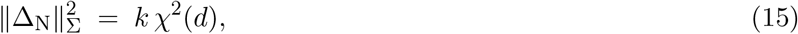

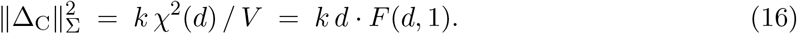

Let *R*_*d*_ = *χ*^2^(*d*)*/d*. By the law of large numbers *R*_*d*_ → 1 in probability, with variance 2*/d*. The Normal step size 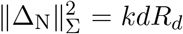 concentrates around *kd*. Every proposal lands on a thin spherical shell of squared Mahalanobis radius approximately *kd*. When *k >* 1 this shell lies outside the typical acceptance region, and there is no variability in the step size to rescue a fraction of proposals.

The Cauchy step size 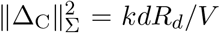 still has *R*_*d*_ → 1, but *V* ∼ *χ*^2^(1) has mean 1 and variance 2 regardless of *d*. The step-size distribution retains a spread over all scales no matter how large *d* gets.

#### Proposition 1 (Dimension-free acceptance for Cauchy)

*Suppose* 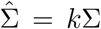 *with k > c >* 0, *and consider the probability that a proposal falls within the squared Mahalanobis radius cd. As d* → ∞:

*(i) Normal:* 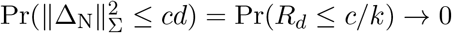 *exponentially in d*.

*(ii) Cauchy:* 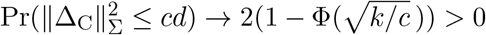 .

#### Proof

For (i), *c/k <* 1 and *R*_*d*_ → 1, so Pr(*R*_*d*_ ≤ *c/k*) decays exponentially by the Cramér bound.

For (ii), rewrite the event as *V* ≥ (*k*/*c*) Rd. Since 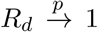 and *V* is independent of *R*_*d*_, Slutsky’s theorem gives

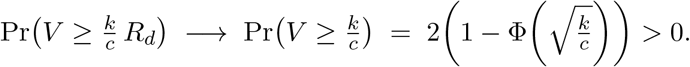

Proposition 1 tells us the Cauchy places mass inside the acceptance region. We can compute the acceptance rate exactly.

Consider a *d*-dimensional product target 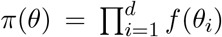 and a random-walk proposal *θ*′= *θ* + *δ*. Roberts et al. (1997) showed that as *d* → ∞, with proposal scale 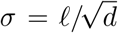, the log-acceptance-ratio

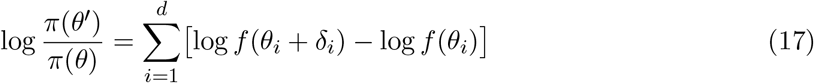

converges by the central limit theorem to a Gaussian. For *f* = *ϕ* (the standard Gaussian density), the log-ratio converges in distribution as *d* → ∞:

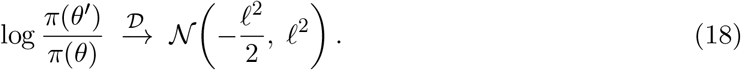

The optimal scale is *l*^*^ ≈ 2.38, giving acceptance rate *a*^*^ ≈ 0.234.

From (18) the Normal acceptance rate is

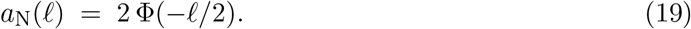

For a Cauchy proposal, *V* ∼ *χ*^2^(1) is drawn once and the effective scale becomes 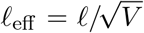 . Conditional on *V* = *v* the log-ratio converges to *N* (−*l*^2^*/*(2*v*), *l*^2^*/v*), so the conditional acceptance rate is 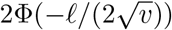 . Averaging over *V* gives

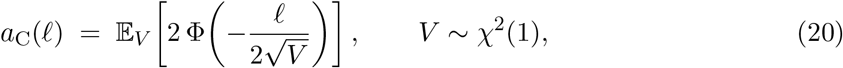

a one-dimensional integral against the *χ*^2^(1) density.

If the proposal is overscaled by a factor *α* relative to optimal, replace *l* by *αl*.

#### Proposition 2 (Robustness gap)

*Under α-fold overscaling (α >* 1*), with l* = *l*^*^ ≈ 2.38:

1. *Normal: a*_N_(*αl*) = 2Φ(−*αl/*2), *decaying as* 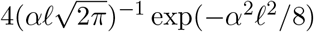 .
2. *Cauchy: a*_C_(*αl*) ≥ 2(1 − Φ(*α*)) · *a*_N_(*l*), positive for all finite *α*.

#### Proof

Part (i) is the standard Gaussian tail bound. For (ii), restrict the expectation in (20) to {*V* ≥ *α*^2^}. On this event 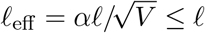, so 2Φ(−*l*_eff_ */*2) ≥ *a*_N_(*l*), and

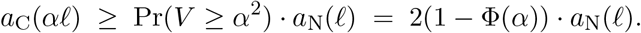

Table 4 spells out representative values. At *α* = 5 the Normal acceptance is 2.6 × 10^−9^ and the Cauchy’s lower bound is 1.3 × 10^−7^, fifty times larger. At *α* = 10 the gap exceeds 10^8^. The crossover is around *α* ≈ 1.44 (*k* ≈ 2): below that the Normal wins because it wastes nothing on extreme scales; above it the Cauchy dominates.

**Table 4:** Acceptance rates under *α*-fold overscaling (*l* = 2.38). The Cauchy column is the lower bound from Proposition 2; the true value is larger.

| $\alpha$ | Normal $a_N(\alpha\ell)$ | Cauchy ≥ | Ratio |
| --- | --- | --- | --- |
| 2 | $1.73 \times 10^{-2}$ | $1.07 \times 10^{-2}$ | 0.62 |
| 3 | $3.57 \times 10^{-4}$ | $6.32 \times 10^{-4}$ | 1.77 |
| 5 | $2.62 \times 10^{-9}$ | $1.34 \times 10^{-7}$ | 50 |
| 10 | $1.18 \times 10^{-32}$ | $3.57 \times 10^{-24}$ | $3.0 \times 10^8$ |

Acceptance rate alone does not capture efficiency; a kernel that takes only tiny steps will accept often but go nowhere. The right measure is the expected squared jump distance (ESJD), given in the diffusion limit by *h*(*ℓ*) = *ℓ*^2^ · *a*(*ℓ*). For the Cauchy,

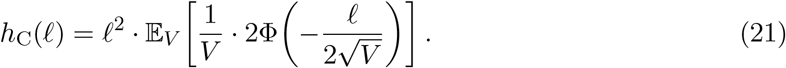

The integral converges despite the 1*/V* factor: for small *V* the acceptance 2Φ(−*ℓ/*( 2√*V*)) ∼ exp(−*ℓ*^2^*/*(8*V*)) kills the integrand. Optimising gives 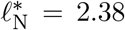 (ESJD 1.33) and 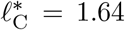 (ESJD 0.71). At optimal tuning the Normal kernel is 1.87 times faster. That is the price of the Cauchy’s robustness: when the proposal covariance is correct, the Cauchy wastes effort on both too-large and too-small steps. Under *α* ≥ 3 overscaling the comparison reverses by orders of magnitude.

### 3.5 The mechanism behind weight concentration

Importance-weighted sequential Monte Carlo carries a well-known difficulty separate from the two failure modes studied above: in high dimensions the importance weights become increasingly uneven, a few particles carrying most of the weight and the effective sample size shrinking accordingly. This weight concentration, whose severe form is the weight degeneracy familiar from the particle-filter literature (Snyder et al., 2008; Bengtsson et al., 2008), is not a failure of either kernel but a property of importance weighting itself, and it affects the Normal and Cauchy kernels alike. As the analysis below shows, it does bear somewhat harder on the Cauchy: the heavy tails that let it escape perturbation overconcentration also place the occasional particle in a novel, low-density region, where it picks up a disproportionately large weight. This section quantifies the mechanism for both kernels.

At each iteration of ABC-SMC, the importance weight assigned to a new particle *θ* is

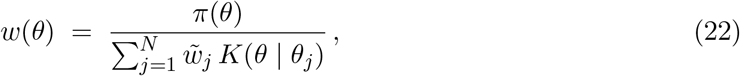

where *π* is the prior, {*θ*_*j*_ } are the previous-iteration particles with normalised weights 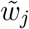, and *K* is the perturbation kernel. A particle that lands far from the previous population has a small denominator (few previous particles contribute kernel density at its location), producing a large weight. This is exactly what happens when the Cauchy kernel’s heavy tails propel a particle into a novel region of the posterior.

To quantify how many previous particles contribute to the denominator, we define the effective number of contributors (ENC):

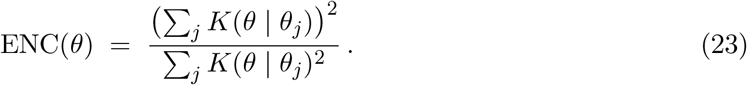

For *N* particles drawn from *N* (0, Σ) with kernel covariance Σ_*K*_ = *c* Σ (where *c* is the kernel coefficient), the inter-particle difference *θ*_*i*_ − *θ*_*j*_ ∼ *N* (0, 2Σ). Setting *r* = 2*/c*, the expected ENC for the Normal kernel has a closed form:

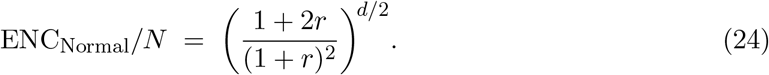

With the standard coefficient *c* = 2 (*r* = 1), this gives (3*/*4)^*d/*2^: exponential decay at rate ≈ 0.86 per dimension. At *d* = 3 roughly 65% of particles contribute; at *d* = 12 only 18%.

The derivation relies on the Gaussian convolution *E*[*K*(Δ)] = *N* (0; 0, Σ_*K*_ +2Σ), which yields *E*[*K*]^2^ and *E*[*K*^2^] in closed form. The ratio *E*[*K*]^2^*/E*[*K*^2^] gives (24) because the normalization constants cancel. The exponent in the Normal kernel density, 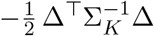, has a coefficient 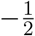 that is independent of *d*.

For the Cauchy kernel (multivariate *t*_1_), the density is proportional to 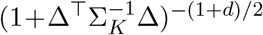. The exponent −(1 + *d*)*/*2 grows with *d*, making the kernel increasingly sensitive to distance differences. No closed-form ENC exists (the Cauchy-Gaussian convolution has no elementary form), but one-dimensional radial quadrature gives exact values (Figure 4b).

**Figure 4:**
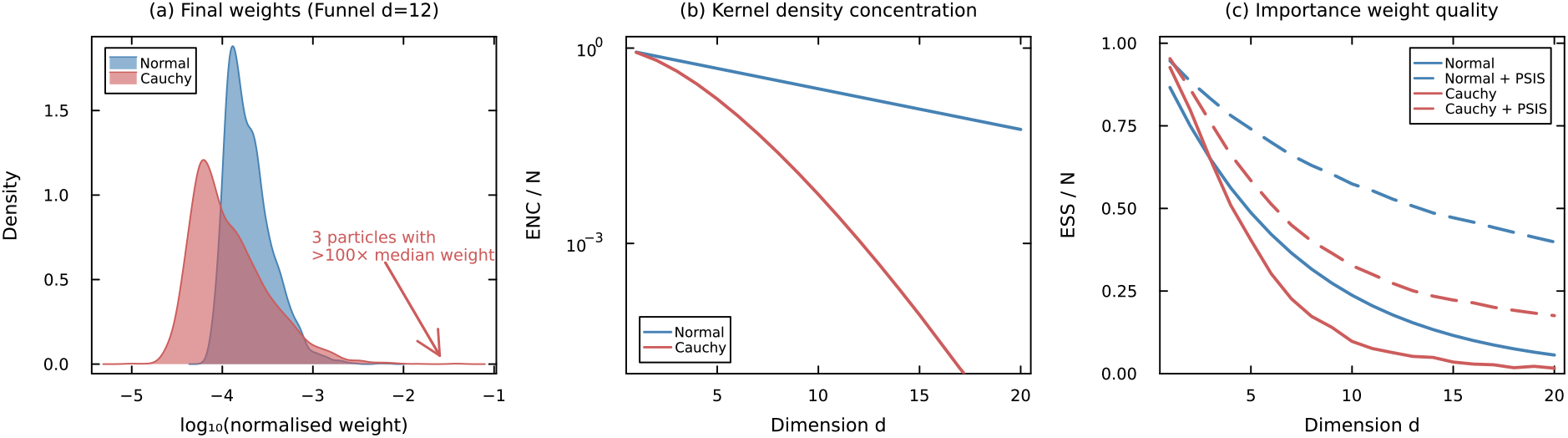
Weight concentration for Normal and Cauchy kernels. (a) Histogram of log_10_-normalised importance weights from the final iteration of ABC-SMC on the Funnel at *d* = 12 (*N* = 4,000, 5×10^7^ simulation budget). The Cauchy weights are more dispersed, reflecting a few high-weight particles that explored regions the previous population did not cover. (b) Effective number of kernel contributors from quadrature: the Normal ENC decays as (3*/*4)^*d/*2^ (equation 24); the Cauchy decays much faster due to its dimension-dependent exponent. (c) ESS/*N* of the importance weights *w* ∝ 1*/* Σ _*j*_ *K*(*θ* | *θ*_*j*_) for *N* = 10,000 particles from *N* (0, *I*_*d*_). Dashed lines show the effect of PSIS post-processing. PSIS substantially recovers ESS/*N* for both kernels, with Normal + PSIS retaining ≈ 0.40 at *d* = 20.

The Cauchy’s dimension-dependent exponent −(1 + *d*)*/*2 makes the kernel density drop off sharply with Mahalanobis distance, so a particle that has moved far from the bulk of the population receives a large weight. The Normal kernel’s constant exponent −1*/*2 produces a flatter density landscape, giving more uniform but less informative weights. The low ESS/*N* of the Cauchy is therefore a natural byproduct of its ability to place particles in regions that the previous population does not cover well.

Pareto-smoothed importance sampling (PSIS; Vehtari et al., 2024) can recover the weight quality without discarding the better-placed particles. PSIS fits a generalised Pareto distribution to the upper tail of the importance weights, taking the largest weights as exceedances above a data-driven threshold. The fitted shape parameter 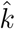 characterises the heaviness of that tail and plays the same role as a classical tail-index estimate. The extreme weights are then replaced by the expected order statistics of the fitted distribution, a monotone transformation that preserves the ranking of the particles and modifies only the upper tail.

Figure 4c shows the effect on both kernels: PSIS raises ESS/*N* substantially for both Normal and Cauchy. At *d* = 20, Normal + PSIS retains ESS/*N* ≈ 0.40 (versus 0.06 without), and Cauchy + PSIS recovers to 0.18 (versus 0.02 without). The kernel coefficient *c* further modulates this: at *d* = 12, increasing *c* from 2 to 8 raises the Cauchy + PSIS ESS/*N* from 0.27 to 0.52. Larger *c* widens the kernel covariance, improving ENC at the cost of lower acceptance rates, but the Cauchy’s heavy tails ensure that exploration is maintained even at high *c*.

Smoothing the weights in this way trades a small bias for a large reduction in variance: the PSIS-adjusted estimator is no longer the unbiased self-normalised importance-sampling estimator, but a regularised one. In the regime where it is needed this trade is favourable, because the raw weights are then heavy-tailed enough (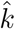 approaching or exceeding one) that the unsmoothed estimator has effectively infinite variance and is itself dominated by a single particle. The shape parameter 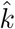 therefore doubles as a diagnostic, with values below roughly 0.7 indicating that the smoothed estimate is reliable (Vehtari et al., 2024). One caveat is specific to the sequential setting: PSIS is justified for a one-shot importance-sampling estimate, whereas here the smoothed weights are applied at every iteration and feed back into the covariance of the next proposal, so we treat the per-iteration use as an empirically validated extension rather than one covered by the existing theory.

### 3.6 The Cauchy perturbation kernel advantage in ABC-SMC grows as over-inflation and overconcentration combine

We ran 10^6^ iterations of both kernels on a *N* (0, *I*_*d*_) target at various *d* and *α*, initialised at stationarity. Table 5 shows the results at *d* = 100, where finite-*d* corrections are small. Every measurable empirical rate sits close to its analytical prediction, lying slightly above it as a finite-*d* effect; the single rate below the 10^−6^ detection limit of 10^6^ draws (the Normal at *α* = 5) is consistent with its far smaller predicted value of 2.7× 10^−9^. At *α* = 3 the Normal acceptance rate has fallen to about 4.7 × 10^−4^ (versus 1.1% for the Cauchy), and by *α* = 5 the Normal accepts no proposals in 10^6^ attempts while the Cauchy still accepts about 0.08%.

**Table 5:** Acceptance rates at *d* = 100: analytical predictions vs empirical random-walk Metropolis.

| $\alpha$ | Normal (theory) | Normal (empirical) | Cauchy (theory) | Cauchy (empirical) |
| --- | --- | --- | --- | --- |
| 1 | 0.234 | 0.237 | 0.164 | 0.167 |
| 1.5 | $7.43 \times 10^{-2}$ | $7.66 \times 10^{-2}$ | $7.97 \times 10^{-2}$ | $8.14 \times 10^{-2}$ |
| 2 | $1.73 \times 10^{-2}$ | $1.92 \times 10^{-2}$ | $3.98 \times 10^{-2}$ | $4.10 \times 10^{-2}$ |
| 3 | $3.57 \times 10^{-4}$ | $4.7 \times 10^{-4}$ | $1.04 \times 10^{-2}$ | $1.12 \times 10^{-2}$ |
| 5 | $2.7 \times 10^{-9}$ | $< 10^{-6}$ | $7.8 \times 10^{-4}$ | $7.9 \times 10^{-4}$ |

**Table 6:** Acceptance rates under controlled overscaling on real posteriors (*d* = 12). Values are averages over 3 replicates × 4 chains. At *k* = 1 the proposal matches the posterior covariance (optimal tuning); at *k >* 1 it is deliberately overscaled. C/N: ratio of Cauchy to Normal acceptance rate. Entries marked “–” had acceptance rate below 10^−4^ for the Normal kernel.

| $k$ | Banana ($d = 12$) | | | Funnel ($d = 12$) | | | Shell ($d = 12$) | | |
| --- | --- | --- | --- | --- | --- | --- | --- | --- | --- |
|  | Normal | Cauchy | C/N | Normal | Cauchy | C/N | Normal | Cauchy | C/N |
| 1 | 0.221 | 0.159 | 0.72 | 0.256 | 0.179 | 0.70 | 0.040 | 0.031 | 0.79 |
| 2 | 0.093 | 0.089 | 0.95 | 0.118 | 0.102 | 0.86 | 0.016 | 0.016 | 0.99 |
| 4 | 0.024 | 0.040 | 1.70 | 0.035 | 0.049 | 1.40 | 0.004 | 0.007 | 1.68 |
| 8 | 0.003 | 0.014 | 4.4 | 0.006 | 0.019 | 3.1 | 0.001 | 0.003 | 5.1 |
| 16 | 0.001 | 0.004 | 5.3 | $< 0.001$ | 0.005 | 8.0 | – | 0.001 | – |
| 32 | $< 0.001$ | 0.001 | 3.1 | – | 0.001 | $\gg 1$ | – | $< 0.001$ | – |

The Gaussian-target verification confirms the formulae, but non-Gaussian posteriors might behave differently. We ran a controlled overscaling experiment on three test problems (Banana, Funnel, Shell) at *d* = 12. For each problem we computed the true posterior covariance Σ from No-U-Turn Sampler (NUTS) reference samples (or analytical samples for the Funnel), then ran both Normal and Cauchy random-walk MH with proposal covariance (2.38^2^*/d*) · *k* · Σ for *k* ∈ {1, 2, 4, 8, 16, 32}. No adaptation was used: the proposal was fixed throughout. Each configuration used 4 chains of 20,000 samples, initialised from reference posterior samples, with 3 independent replicates.

At *k* = 1 the Normal wins by a factor of roughly 1.4, consistent with the optimal-tuning ESJD ratio of 1.87. The crossover occurs between *k* = 2 and *k* = 4 for all three problems, matching the analytical prediction of *α* ≈ 1.44 (*k* ≈ 2). At *k* = 8 the Cauchy accepts 3 to 5 times more proposals; at *k* = 16 the advantage reaches 5 to 8 times where the Normal still accepts at all. These three posteriors have very different geometries: Banana has a curved ridge, Funnel has scales that vary by orders of magnitude across directions, and Shell has an annular structure with no density at the mode. Yet the crossover point (*k* = 2–4) and the scaling of the Cauchy advantage are the same across all three, matching the problem-agnostic prediction from Gaussian-target theory. This is because the theory depends only on properties of the proposal distribution (concentration of *χ*^2^(*d*)*/d*), not on the target.

The MH analysis isolates the proposal mechanism and confirms the Cauchy advantage under overscaling. We now return to the original ABC-SMC setting to check that this advantage survives when importance weights are present.

We ran both kernels (correctly specified, so the importance weights use the same kernel density as the proposal) on all five test problems at *d* = 3 and *d* = 12, with *N* = 4,000 particles (5 replicates for synthetic problems, 3 for Gene). Posterior quality was measured by the sliced Wasserstein distance (SWD; Section 2) and three variants of the classifier two-sample test (C2ST), each comparing resampled ABC posterior samples against reference samples: the joint C2ST (classifier on all *d* dimensions simultaneously), the marginal C2ST (mC2ST, average of *d* one-dimensional classifiers), and the pairwise C2ST (pC2ST, average of all 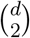 two-dimensional classifiers). For C2ST, 0.5 is perfect posterior recovery and 1.0 is complete failure; for SWD, lower is better. We also report the effective sample size as a fraction of *N* (ESS/*N*) and the final overscaling ratio 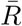.

Table 7 summarises the results. At *d* = 3 the two kernels are essentially indistinguishable on all five problems and all metrics, with differences at the third decimal of SWD that are smaller than the run-to-run variance. At *d* = 12 the picture is more nuanced: each problem occupies a different regime of the overinflation–concentration interaction.

**Table 7:** Normal vs Cauchy kernel in ABC-SMC across five problems (*N* = 4,000, 10^8^ simulation budget, *p*_acc,min_ = 10^−4^; 5 replicates for synthetic problems, 3 for Gene). SWD: sliced Wasserstein distance (1000 random projections; lower = closer to reference posterior). C2ST variants: joint (all dimensions), marginal (per-dimension average), pairwise (all 2D pairs); 0.5 is perfect, 1.0 is failure. ESS/*N* : effective sample size as a fraction of *N* . 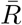 : mean overscaling ratio of the kernel covariance relative to the true posterior covariance. Reference posteriors are analytical (Gaussian, Funnel) or from NUTS (Banana, Shell, Gene).

| Problem | Kernel | Posterior quality (↓ better) | | | | Iters | Weights | | Sims |
| --- | --- | --- | --- | --- | --- | --- | --- | --- | --- |
| | | SWD | Joint | Marg. | Pair. | | ESS/$N$ | $\bar{R}$ | |
| $d = 3$ | | | | | | | | | |
| Gaussian | Normal | 0.005 | 0.51 | 0.50 | 0.51 | 26 | 0.78 | 2 | 92.7M |
|  | Cauchy | 0.008 | 0.57 | 0.51 | 0.53 | 26 | 0.25 | 2 | 97.8M |
| Funnel | Normal | 0.005 | 0.53 | 0.51 | 0.52 | 28 | 0.63 | 2 | 94.2M |
|  | Cauchy | 0.006 | 0.53 | 0.51 | 0.52 | 28 | 0.57 | 2 | 98.0M |
| Banana | Normal | 0.030 | 0.63 | 0.55 | 0.58 | 39 | 0.52 | 2 | 94.4M |
|  | Cauchy | 0.034 | 0.64 | 0.55 | 0.58 | 39 | 0.14 | 2 | 94.9M |
| Shell | Normal | 0.041 | 0.50 | 0.50 | 0.51 | 19 | 0.80 | 2 | 100M |
|  | Cauchy | 0.048 | 0.52 | 0.50 | 0.50 | 19 | 0.50 | 2 | 100M |
| Gene | Normal | 0.124 | 0.82 | 0.54 | 0.59 | 26 | 0.71 | 2 | 100M |
|  | Cauchy | 0.102 | 0.80 | 0.53 | 0.56 | 28 | 0.45 | 1 | 100M |
| $d = 12$ | | | | | | | | | |
| Gaussian | Normal | 0.034 | 0.88 | 0.56 | 0.60 | 53 | 0.20 | 3 | 97.7M |
|  | Cauchy | 0.031 | 0.85 | 0.57 | 0.60 | 52 | 0.06 | 3 | 93.6M |
| Funnel | Normal | 0.122 | 0.98 | 0.60 | 0.68 | 57 | 0.09 | 100 | 91.9M |
|  | Cauchy | 0.061 | 0.96 | 0.59 | 0.66 | 58 | 0.06 | 17 | 84.8M |
| Banana | Normal | 0.903 | 1.00 | 0.99 | 1.00 | 64 | 0.39 | 3,737 | 73.7M |
|  | Cauchy | 0.018 | 0.83 | 0.53 | 0.56 | 109 | 0.14 | 2 | 73.7M |
| Shell | Normal | 0.039 | 0.79 | 0.51 | 0.52 | 41 | 0.22 | 2 | 100M |
|  | Cauchy | 0.029 | 0.64 | 0.50 | 0.51 | 40 | 0.48 | 2 | 100M |
| Gene | Normal | 0.687 | 1.00 | 0.70 | 0.86 | 33 | 0.44 | 277 | 100M |
|  | Cauchy | 0.059 | 1.00 | 0.58 | 0.73 | 91 | 0.23 | 0.3 | 100M |

The Gaussian model at *d* = 12 is instructive. Although the sample mean 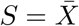 is a sufficient statistic in the Bayesian sense, the ABC distance ∥*S*_sim_ − *S*_obs_∥ operates in ℝ^*d*^, and at *d* = 12 the acceptance ball is large enough relative to the posterior that overinflation is non-trivial. The Cauchy kernel achieves a modest improvement in SWD (0.031 vs 0.034) and joint C2ST (0.85 vs 0.88), though the marginal metrics are similar.

The five problems at *d* = 12 illustrate distinct regimes of the overinflation–concentration interaction, and the magnitude of the Cauchy advantage tracks the degree of overinflation across all of them.

The Funnel 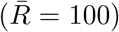 shows the Cauchy halving SWD (from 0.122 to 0.061) and reducing 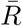 from 100 to 17, with fewer total simulations (84.8M vs 91.9M).

The Banana is the most dramatic case. With the longer budget, the Cauchy kernel achieves a 50 × reduction in SWD (from 0.903 to 0.018), bringing mC2ST from 0.99 (complete failure) to 0.53 (near-perfect recovery), and reducing 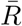 from 3,737 to 2. The Cauchy runs 109 iterations vs the Normal’s 64 within a similar simulation budget (∼74M), demonstrating the virtuous cycle: by maintaining acceptance where the Normal stalls, it pushes to tighter tolerances until the overinflation resolves itself entirely.

Shell shows both kernels performing well with the corrected radii-only summary statistics 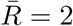, but the Cauchy still achieves a meaningful advantage at *d* = 12: SWD 0.029 vs 0.039 and joint C2ST 0.64 vs 0.79. Shell is also the only problem where the Cauchy achieves higher ESS/*N* than the Normal (0.48 vs 0.22). On all other problems the Normal’s concentrated step sizes produce more uniform weights, but on the annular posterior the Cauchy’s variable step sizes spread proposals across different radii, and those that land near the shell receive good weights, yielding a less degenerate particle population.

The gene expression problem confirms that the Cauchy advantage extends to a realistic biological model. The hierarchical structure creates a funnel-shaped posterior in which the spread parameter *τ* controls the effective dimensionality of the random effects *ε*_*i*_. At *d* = 12 the Normal kernel is visibly struggling: 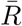 inflates to 277, the acceptance rate drops, and the algorithm stalls at 33 iterations against the 10^8^ budget with SWD = 0.69. The Cauchy kernel maintains 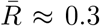, sustains acceptance through 91 iterations within the same budget, and achieves a 12 × reduction in SWD (0.059 vs 0.687). Unlike the other problems, the gene model’s 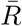 sits below 1, meaning the particle cloud is narrower than the reference posterior rather than wider. This is expected: the gene model is the only problem with a deterministic ABC distance, so there is no irreducible simulator-noise floor, and the ABC target contracts toward the moment-matching solution as the tolerance falls, leaving a converged cloud narrower than the full-likelihood NUTS reference. The result is benign under-dispersion, the converse of the covariance overinflation behind the Normal kernel’s failure; for this model 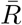 partly reflects how far the run has progressed, since each additional iteration contracts the cloud further against the deterministic distance.

The marginal C2ST corroborates the gap (0.58 for the Cauchy versus 0.70 for the Normal) and the pairwise C2ST widens it further (0.73 vs 0.86). With the reduced 5 × 10^7^ budget (Table 8), the Cauchy kernel already achieves SWD = 0.066 and 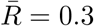, essentially the full 10^8^ result at half the cost, so Gene is also the problem where the budget reduction has the smallest effect on the Cauchy.

**Table 8:**
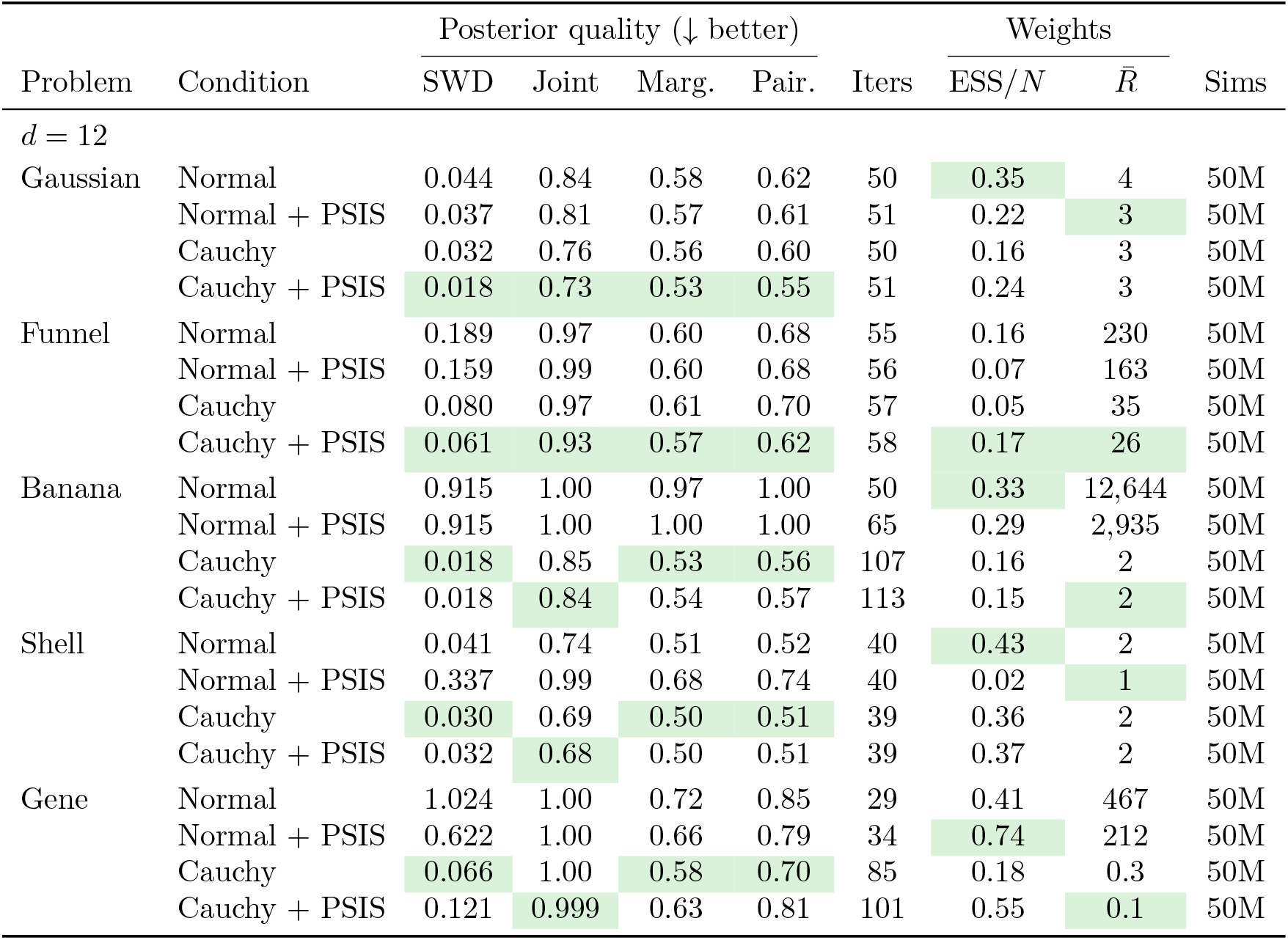
Fixed-budget comparison (*N* = 4,000, 5 × 10^7^ simulations, no *p*_acc,min_ stopping, 3 replicates). +PSIS applies Pareto-smoothed importance sampling to the weights after each iteration.

| Problem | Condition | Posterior quality (↓ better) | | | | Iters | Weights | | Sims |
| --- | --- | --- | --- | --- | --- | --- | --- | --- | --- |
| | | SWD | Joint | Marg. | Pair. | | ESS/$N$ | $\bar{R}$ | |
| $d = 12$ | | | | | | | | | |
| Gaussian | Normal | 0.044 | 0.84 | 0.58 | 0.62 | 50 | 0.35 | 4 | 50M |
|  | Normal + PSIS | 0.037 | 0.81 | 0.57 | 0.61 | 51 | 0.22 | 3 | 50M |
|  | Cauchy | 0.032 | 0.76 | 0.56 | 0.60 | 50 | 0.16 | 3 | 50M |
|  | Cauchy + PSIS | 0.018 | 0.73 | 0.53 | 0.55 | 51 | 0.24 | 3 | 50M |
| Funnel | Normal | 0.189 | 0.97 | 0.60 | 0.68 | 55 | 0.16 | 230 | 50M |
|  | Normal + PSIS | 0.159 | 0.99 | 0.60 | 0.68 | 56 | 0.07 | 163 | 50M |
|  | Cauchy | 0.080 | 0.97 | 0.61 | 0.70 | 57 | 0.05 | 35 | 50M |
|  | Cauchy + PSIS | 0.061 | 0.93 | 0.57 | 0.62 | 58 | 0.17 | 26 | 50M |
| Banana | Normal | 0.915 | 1.00 | 0.97 | 1.00 | 50 | 0.33 | 12,644 | 50M |
|  | Normal + PSIS | 0.915 | 1.00 | 1.00 | 1.00 | 65 | 0.29 | 2,935 | 50M |
|  | Cauchy | 0.018 | 0.85 | 0.53 | 0.56 | 107 | 0.16 | 2 | 50M |
|  | Cauchy + PSIS | 0.018 | 0.84 | 0.54 | 0.57 | 113 | 0.15 | 2 | 50M |
| Shell | Normal | 0.041 | 0.74 | 0.51 | 0.52 | 40 | 0.43 | 2 | 50M |
|  | Normal + PSIS | 0.337 | 0.99 | 0.68 | 0.74 | 40 | 0.02 | 1 | 50M |
|  | Cauchy | 0.030 | 0.69 | 0.50 | 0.51 | 39 | 0.36 | 2 | 50M |
|  | Cauchy + PSIS | 0.032 | 0.68 | 0.50 | 0.51 | 39 | 0.37 | 2 | 50M |
| Gene | Normal | 1.024 | 1.00 | 0.72 | 0.85 | 29 | 0.41 | 467 | 50M |
|  | Normal + PSIS | 0.622 | 1.00 | 0.66 | 0.79 | 34 | 0.74 | 212 | 50M |
|  | Cauchy | 0.066 | 1.00 | 0.58 | 0.70 | 85 | 0.18 | 0.3 | 50M |
|  | Cauchy + PSIS | 0.121 | 0.999 | 0.63 | 0.81 | 101 | 0.55 | 0.1 | 50M |

Figure 5 shows how the kernel comparison scales with the number of genes, from *d* = 3 (6 parameters) to *d* = 20 (23 parameters), with a uniform simulation budget of 10^8^ at every dimension so that each configuration is given the same resource. At *d* ≤9 all four configurations (Normal, Cauchy, Normal + PSIS, Cauchy + PSIS) achieve SWD below 0.2 and 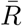 at most 2; the kernels are indistinguishable on posterior quality and PSIS contributes nothing because the particle weights are already healthy. The first departure appears at *d* = 12: the Normal kernel stalls at 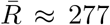 and SWD ≈ 0.69, while both Cauchy configurations and Normal + PSIS remain at SWD ≤ 0.12 and 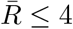. At *d* = 15 both vanilla and PSIS-smoothed Normal kernels fail (SWD ≈ 1.07 and 0.86 respectively,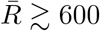), while the Cauchy variants stay at SWD ≤ 0.12 and 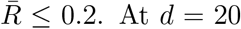 raw Cauchy also fails, though through a different mechanism: weight degeneracy (ESS/*N* ≈ 0.03) produces high run-to-run variance in SWD (individual replicates at 0.93, 2.33, 4.49). Only Cauchy + PSIS survives at *d* = 20, achieving SWD = 0.128 and 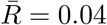 with ESS/*N* = 0.37 across 145 iterations. The two techniques address complementary failure modes: the Cauchy kernel neutralises perturbation overconcentration, and PSIS controls weight degeneracy, with the threshold at which each technique becomes necessary lying at a different dimension.

**Figure 5:**
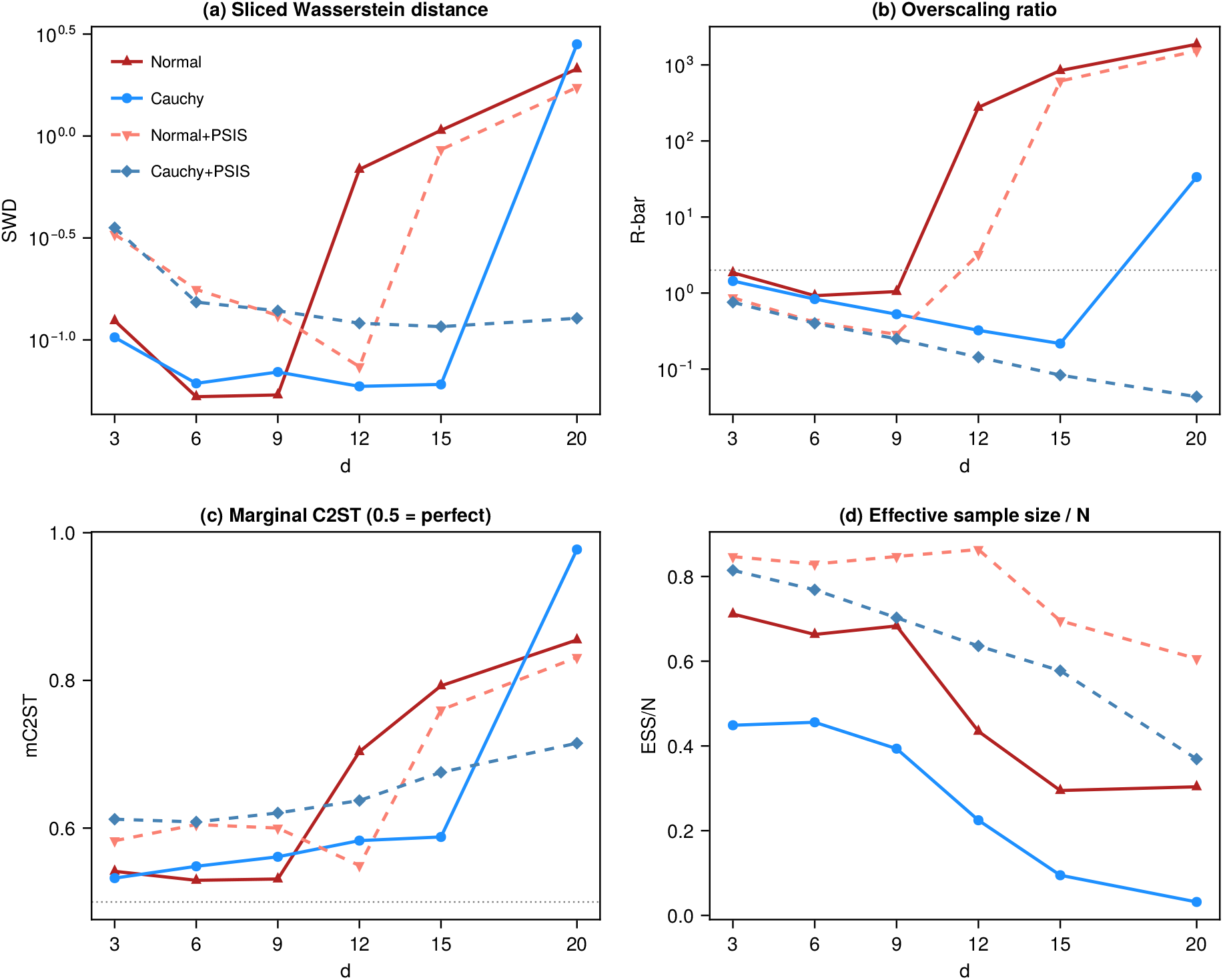
Dimension scaling on the gene expression problem (*N* = 4,000, *p*_acc,min_ = 10^−4^, flat 10^8^ simulation budget at every *d*, 3 replicates). Both kernels perform well up to *d* = 9. The Normal kernel starts to struggle at *d* = 12 and fails at *d* = 15; Normal + PSIS extends the Normal kernel’s reach by one dimension but also fails at *d* = 15. Cauchy and Cauchy + PSIS both survive through *d* = 15; only Cauchy + PSIS survives at *d* = 20, where raw Cauchy encounters weight degeneracy (ESS/*N* ≈ 0.03, with individual replicate SWDs ranging from 0.93 to 4.49).

The sliced Wasserstein distance provides the clearest picture of the Cauchy advantage at *d* = 12, where the joint C2ST often saturates at 1.0 and cannot distinguish the two kernels. The Cauchy reduces SWD on all five problems, with the improvement ranging from modest (1.1× on Gaussian) to dramatic (12× on Gene, 50× on Banana), confirming that the improvement is distributional, not merely a point-estimate artefact. The marginal and pairwise C2ST variants corroborate this: the marginal C2ST has the best dynamic range among the classifier-based metrics, while the pairwise C2ST captures correlation mismatches that marginals miss.

## 4 Discussion

Two independent failure modes afflict ABC-SMC with Normal perturbation kernels. Covariance overinflation, driven by information loss in the summary statistics, causes the kernel covariance to overshoot the true posterior covariance by a large factor 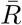 . Perturbation overconcentration, driven by dimension via sd 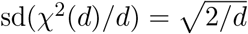, forces every Normal proposal onto a thin shell at the overscaled radius. Neither alone is fatal. Overinflation without concentration is harmless: at *d* = 3 with 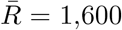 both kernels perform identically. Concentration without overinflation is harmless: with sufficient statistics 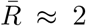 and the Normal kernel works well. Only the combination is lethal, and only for the Normal kernel.

The Cauchy kernel survives because its step size includes a factor 1*/V* with *V* ∼ *χ*^2^(1), which never concentrates regardless of *d*. This mechanism is problem-agnostic: the acceptance rate formulae, the crossover at *α* ≈ 1.44 (*k* ≈ 2), and the ESJD comparison all follow from properties of the proposal distribution and do not depend on the target. Controlled experiments on three non-Gaussian posteriors confirm the predicted crossover, and full ABC-SMC runs show the Cauchy substantially reducing the sliced Wasserstein distance to the reference posterior across all five test problems at *d* = 12, including a biologically motivated gene expression model.

Beyond tolerating overinflation, the Cauchy kernel also reduces it. Table 7 shows that 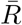 is lower under the Cauchy at *d* = 12 on three of five problems: 17 vs 100 (Funnel), 2 vs 3,737 (Banana), and 0.3 vs 277 (Gene), with the Gaussian and Shell comparable at 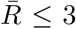 for both kernels. The likely mechanism is that higher acceptance rates allow the Cauchy to push to tighter tolerances (more iterations within the same simulation budget), producing a particle cloud that is closer to the true posterior and therefore a less overinflated kernel covariance at the next iteration. This creates a virtuous cycle: better proposals lead to tighter tolerances, which lead to less overinflation, which further improves proposal quality.

A further source of waste under overinflated kernels is prior rejection. When the prior has bounded support (as is typical in ABC applications), proposals from an overinflated kernel frequently land outside the prior bounds and are discarded before the simulator is ever called. This means that much of the computational budget is consumed by proposals that cannot possibly contribute to the population. Which kernel suffers more from prior rejection depends on the regime. The Cauchy’s heavy tails cut both ways: they keep some proposals short even when the covariance is overinflated, but they also send some proposals far past the prior bounds. On Shell at 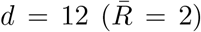 with a fixed 50M simulation budget, the Normal kernel issued 52M proposals over 40 iterations and the Cauchy 58M over 39, so when overinflation is mild the Cauchy pays a slightly larger out-of-bounds overhead. The Normal kernel’s disadvantage emerges only under severe overinflation, where its concentrated proposal shell lies almost entirely outside the prior and nearly every proposal is rejected before simulation, while the Cauchy retains a usable in-support fraction.

The decomposition suggests two complementary strategies for improving ABC-SMC. The first is to use Cauchy kernels, which neutralises overconcentration at the cost of a 1.87× efficiency penalty when the proposal is correctly scaled. The second is to improve the summary statistics, which reduces overinflation directly. The two are not substitutes: better summaries help at any dimension, and the Cauchy helps at any level of overinflation provided *d* is large enough for concentration to matter. A third strategy would target the covariance estimate itself, through shrinkage or locally adapted proposals (Filippi et al., 2013). This is complementary rather than competing: such estimators can lower 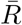 but cannot push it below the structural floor set by the excess width of the ABC posterior, and they add estimation cost and tuning that the Cauchy avoids. The Cauchy is attractive precisely because it tolerates whatever overinflation remains at no additional cost.

Since overinflation is hard to predict in advance and the Cauchy’s worst-case penalty is bounded at 1.87×, the Cauchy remains a safe default. But understanding that the root cause is information loss in the summaries, not dimension itself, raises the question of whether over-inflation can be avoided by using better summary statistics. In principle, yes: with sufficient statistics, the analytical model shows 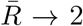 at any dimension, and the Normal kernel works normally. In practice, no: ABC is used precisely for models where the likelihood is intractable, and these are almost invariably models where no finite-dimensional sufficient statistic exists (sufficient statistics require exponential family structure). The summary statistics are necessarily insufficient, and the resulting overinflation is a fundamental feature of the method, not a fixable deficiency.

Increasing the particle count *N* does not close the overinflation gap. *N* controls how well the particle cloud approximates the ABC posterior, not how close the ABC posterior is to the true posterior, and the 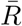 vs *N* experiment confirmed this: 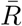 is flat at roughly 880 across *N* = 500 to 8,000 on the Funnel at *d* = 12. The companion experiment with the Cauchy kernel (Appendix B) shows the same *N* -independence: Cauchy 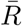 on the Funnel at *d* = 12 is also flat across *N*, at a lower value (≈ 630 vs the Normal’s ≈ 880) but with the same absence of a sample-size trend. Overscaling is structural for both kernels, and *N* controls only the sampling error around the floor, not its location.

The Cauchy kernel is therefore not a workaround for a temporary limitation but a response to a structural one. Overinflation is baked into ABC-SMC by the information loss in the summary statistics, and this information loss is inherent to the class of problems that require ABC. The only remaining question is whether the dimension is high enough for concentration to make overinflation lethal, and the answer is typically yes once *d* exceeds about 5.

This structural reading has a natural home in the theory of adaptive multilevel splitting: adaptive-tolerance ABC-SMC is itself such a scheme, in which a particle population is driven through a sequence of levels set adaptively as empirical quantiles of the current ensemble. Seen this way, the overinflation we document is a property of the adaptive construction rather than of the particle count: the population approximates the ABC posterior at the current tolerance, which is wider than the true posterior whenever the tolerance is positive, and this gap does not shrink as *N* grows (Table 1). For schemes of exactly this kind, the fluctuation analysis of Cérou and Guyader (2016) gives a central limit theorem for the splitting estimator under adaptively chosen levels, quantifying the asymptotic variance contributed by the data-driven thresholds. That analysis treats the adaptive levels while assuming a well-behaved mutation step, whereas we study the mutation step itself, the perturbation kernel. The two are therefore complementary, accounting respectively for the cost of adaptive thresholds and for the cost of an overscaled kernel, which the Cauchy attenuates.

The Cauchy advantage is specific to algorithms that cannot adapt their covariance to the true target. In standard adaptive MH the chain samples from the true posterior, the covariance estimator converges to the true posterior covariance, and the proposal is correctly scaled (*k* ≈ 1). At correct scaling the Normal kernel is 1.87× faster in ESJD, and the Cauchy’s heavy tails waste proposals on steps that are either far too large or far too small. Other particle methods with structural covariance mismatch, such as sequential Monte Carlo samplers with likelihood-tempered sequences, may also benefit from heavy-tailed kernels.

Convergence guarantees are not affected by the switch to Cauchy proposals. Geometric ergodicity of random-walk Metropolis requires exponentially light tails on the target (Mengersen and Tweedie, 1996; Jarner and Hansen, 2000), not on the proposal, and in ABC the target is bounded by the prior support. The heavy-tailed Cauchy density in the importance-weight denominator does not produce infinite-variance weights either: when the prior restricts parameters to a compact set, the Cauchy density is bounded away from zero on that set, and the weights stay bounded.

The proposal mechanism and the importance weights can be optimised independently. The Cauchy addresses overinflation on the proposal side, but its heavy tails concentrate the importance weights (lower ESS/*N*). We investigated per-iteration PSIS weight smoothing (Vehtari et al., 2024) as a remedy: Table 8 shows the effect of applying PSIS to the importance weights after each ABC-SMC iteration. The interaction between PSIS and kernel shape is striking. For the Cauchy kernel, PSIS improves posterior quality when there is residual overinflation for it to fix: on Gaussian it reduces SWD from 0.032 to 0.018, and on Funnel from 0.080 to 0.061. When the Cauchy kernel already achieves 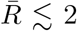 without PSIS (Banana, Shell, Gene at the 5 × 10^7^ budget), PSIS is neutral: Banana is unchanged at SWD = 0.018, Shell shifts from 0.030 to 0.032, and Gene shifts from 0.066 to 0.121, all still well within the distributional-recovery regime. For the Normal kernel, per-iteration PSIS can be actively harmful: on Shell it increases SWD from

0.041 to 0.337 (with one replicate collapsing entirely to 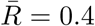), and on Funnel and Gaussian it produces worse or comparable results. The mechanism is that PSIS trims the largest weights, which for the Normal kernel’s concentrated proposals are often the only particles near the target; smoothing them down destroys the covariance signal for the next iteration. The Cauchy’s more variable step sizes produce a broader weight distribution that is robust to this smoothing. The upshot is a cleaner rule than the original claim that PSIS is uniformly beneficial: PSIS helps the Cauchy kernel only when the weight-distribution pathology is the remaining bottleneck, which in practice means the problems where raw Cauchy leaves 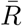 at tens rather than units. On the five *d* = 12 problems studied here that is Gaussian and Funnel; PSIS is also essential at *d* = 20 on the gene model (Figure 5), where raw Cauchy develops weight degeneracy (ESS/*N* ≈ 0.03) and only Cauchy + PSIS produces a posterior close to the reference.

## A Fixed-budget comparison

To control for the possibility that the Cauchy advantage in Table 7 arises from consuming more simulations before hitting *p*_acc,min_, Table 8 repeats the comparison with a fixed budget of 5 × 10^7^ simulations and no acceptance-rate stopping criterion (*p*_acc,min_ = 0). The +PSIS conditions apply Pareto-smoothed importance sampling to the weights after each iteration, modifying the covariance estimate used for the next proposal.

## B Particle count does not close the overscaling gap for any configuration

Table 1 showed that for the Normal kernel on the Funnel at *d* = 12, the final overscaling ratio 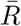 is flat across *N* ∈ [500, 8,000]. This is consistent with the analytical derivation in Section 3.3, which shows 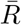 as a property of the ABC posterior *π*_*ε*_ rather than of the particle cloud used to approximate it. A natural question is whether the same *N* -independence holds for the other three configurations, or whether the Cauchy’s lower 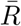 in Table 7 or the behaviour of the PSIS variants reflect finite-sample effects that would change with more particles.

Table 9 answers this question by repeating the Table 1 experiment for all four configurations: same problem (Funnel *d* = 12), same stopping criterion (*p*_acc,min_ = 0.01, 10^8^ simulation budget), same *N* values, three replicates per row.

**Table 9:** Companion to Table 1: final overscaling ratio 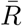 and effective sample size across particle counts for all four configurations on the Funnel at *d* = 12 (*p*_acc,min_ = 0.01, 10^8^ sim budget, 3 replicates per row; reported values are 3-rep means). 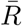 is flat with respect to *N* for every configuration. PSIS has little effect on 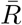 under either kernel but substantially increases ESS/*N*, particularly for the Cauchy.

| Configuration | $N$ | Iters | $\bar{R}$ | Total sims | $\text{ESS}/N$ |
| --- | --- | --- | --- | --- | --- |
| Normal | 500 | 40 | 852 | 1.8M | 81% |
|  | 1,000 | 40 | 863 | 3.6M | 78% |
|  | 2,000 | 39 | 872 | 7.0M | 81% |
|  | 4,000 | 40 | 878 | 14.1M | 82% |
|  | 8,000 | 40 | 880 | 28.5M | 81% |
| Normal + PSIS | 500 | 42 | 855 | 1.9M | 92% |
|  | 1,000 | 42 | 875 | 3.8M | 89% |
|  | 2,000 | 41 | 855 | 7.3M | 89% |
|  | 4,000 | 40 | 865 | 14.4M | 88% |
|  | 8,000 | 41 | 868 | 29.2M | 87% |
| Cauchy | 500 | 47 | 599 | 1.7M | 12% |
|  | 1,000 | 46 | 631 | 3.2M | 19% |
|  | 2,000 | 47 | 587 | 6.5M | 14% |
|  | 4,000 | 46 | 630 | 12.8M | 19% |
|  | 8,000 | 46 | 659 | 25.2M | 22% |
| Cauchy + PSIS | 500 | 50 | 646 | 1.8M | 56% |
|  | 1,000 | 50 | 576 | 3.6M | 53% |
|  | 2,000 | 49 | 590 | 7.0M | 48% |
|  | 4,000 | 48 | 613 | 13.6M | 49% |
|  | 8,000 | 48 | 611 | 26.9M | 45% |

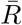 is flat across *N* for every configuration: Normal at ≈ 870, Normal + PSIS at ≈ 864, Cauchy at ≈ 621, Cauchy + PSIS at ≈ 607, each with variation under 10% across a 16× range of *N* and no monotone trend. The 16× increase in particle count does not change the overscaling floor for any kernel–weighting combination. This extends the analytical result of Section 3.3 from the Gaussian model with sufficient statistics to an empirical setting with insufficient summary statistics, and confirms that structural overscaling is a property of the kernel and the stopping criterion rather than of the particle cloud approximating the ABC posterior.

PSIS has essentially no effect on 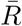 under either kernel at this stopping criterion, with Normal vs Normal + PSIS differing by under 1% in the typical 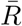 and Cauchy vs Cauchy + PSIS differing by roughly 2%. The mechanism is that 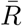 is determined by the weighted empirical covariance at the final iteration, and PSIS smooths only the extreme weights; when the weight distribution is not heavy-tailed (Normal kernel) or when the stopping criterion is loose (both kernels at *p*_acc,min_ = 0.01), the smoothing does not move the bulk of the covariance estimate appreciably. What PSIS does change is the effective sample size fraction: Normal + PSIS raises ESS/*N* from roughly 0.80 to 0.89, and Cauchy + PSIS raises it from roughly 0.17 to 0.51, a threefold improvement. This matters for any downstream inference that uses the weighted particle cloud directly, such as posterior means, credible intervals, or covariance estimates, even though it does not change the overscaling floor.

The Cauchy floor at 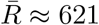 is about 71% of the Normal floor at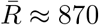, and the Cauchy variants terminate at a slightly higher iteration count (46–50 vs 40–42) under *p*_acc,min_ = 0.01. Under this loose stopping criterion the four configurations produce broadly similar iteration counts and similar particle clouds, so the Cauchy’s modest 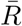 advantage here reflects the direct effect of the perturbation shape on the covariance estimator rather than the virtuous cycle described in Section 3.2: the Cauchy’s variable step sizes produce a slightly less anisotropic particle population at termination, but without the many additional iterations that a tighter *p*_acc,min_ would allow, the full gap seen in Table 7 (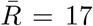 for Cauchy versus 100 for Normal at *p*_acc,min_ = 10^−4^) does not materialise. The large Cauchy advantage is therefore not a finite-*N* artefact: it is a dynamical effect of letting the Cauchy run further before the algorithm terminates.

